# Data-driven prioritization of mouse strains for improved preclinical modeling of rare and common disease

**DOI:** 10.64898/2026.04.27.721175

**Authors:** Robyn L. Ball, Alyssa Klein, Matthew W. Gerring, Alexander K. Berger-Liedtka, Matthew J. Kim, Melissa L. Berry, Michael A. Gargano, Gaurab Mukherjee, Heidi S. Fisher, Tessa Nichols-Meade, Francisco Castellanos, Cynthia L. Smith, Guy Karlebach, Stephen A. Murray, Carol J. Bult, Peter N. Robinson, Elissa J. Chesler

## Abstract

Choosing an appropriate mouse genetic background is a persistent challenge for successful translation of preclinical disease modeling. We present Strain Recommender, a genomic framework that prioritizes inbred mouse strains as relatively vulnerable or resilient to a disease state using disease-associated gene signatures and strain-specific transcriptome predictions. The method represents disease states as weighted gene scores, ranks 657 strains based on resemblance to the disease state, and estimates uncertainty via a permutation-derived false positive rate (FPR). In a prospective validation of connective tissue disorder predictions, vulnerable and resilient Collaborative Cross strains showed significantly different cardiovascular abnormalities. In a global retrospective validation predicting previously reported strain background effects, Strain Recommender achieved ≥ 90% sensitivity for 86.6% of diseases with 94.4% mean sensitivity (95% CI: 94.0-94.8%) across 5,890 diseases, including 92.3% (95% CI: 91.6-93.0%) for 2,598 rare diseases, demonstrating its potential to improve the validity of mouse models of human disease.

---

Model organisms provide a powerful platform for biomedical research, enabling controlled experiments to test disease mechanisms, characterize the effect of genetic variation and assess therapeutic strategies. Inbred mouse strains, the genetic backgrounds on which disease models are built are highly genetically and phenotypically diverse (Beck et al., 2000). They share roughly 90% of disease-related genes with humans (Mayor, 2002), and their fixed genomes support reproducibility and longitudinal studies. Moreover, valid mouse models of disease recapitulate human disease states (Robinson et al., 2013; Pan et al., 2023; Rydell-Törmänen and Johnson, 2019) and have accelerated the development of treatments and interventions (Domínguez-Oliva et al., 2023; Singh and Seed, 2021), particularly for conditions in which *in vivo* assays are indispensable.

Despite their widespread use, the translational success of mouse disease models can be limited by the choice of strain background. The choice of strains is largely based on convention rather than systematic optimization and is often constrained to a small number of deeply characterized strains, such as C57BL/6 (Mouse Genome Sequencing Consortium, 2002) due to historical precedent (Silva et al., 1997), ease of use, and resource availability. For Mendelian disorders, the relevance of a ‘mutant mouse’ to human disease states depends not only on the gene mutation itself but critically on the genetic background in which it is expressed (Rivera and Tessarollo, 2008), with the same mutation or treatment producing markedly different phenotypic outcomes across strains (Justice and Dhillon, 2016; Sittig et al., 2016; Coley et al., 2015). For complex diseases, susceptibility is shaped by polygenic interactions in the biological network, which vary substantially among inbred strains. A strain chosen without regard to this genomic context can compromise translational relevance, yielding models that incompletely reflect human disease phenotypes (Loewa et al, 2023; Mak et al., 2014; Ineichen et al, 2024). Systematic, evidence-based strain selection thus represents an unmet need in preclinical research design.

Hundreds of inbred strains (and hundreds of thousands of potential hybrids) exist, many of which may better capture disease susceptibility—or resilience—than the most widely used laboratory strains (Lutz and Osborne, 2013; Sellers 2017). However, background effects are difficult to predict in advance and comprehensive experimental screening across backgrounds is rarely feasible: engineering, breeding, and phenotyping multiple models is costly, time-consuming and increases animal use. As a result, investigators often commit to a background early, without quantitative evidence of alignment with the disease state they aim to model (Dumont et al., 2024, Festing, 1976), yet identifying the optimal strain *a priori* would minimize unnecessary animal use and improve the likelihood of translational success.

To address this challenge, we developed ‘Strain Recommender’, a data-driven genomic approach to rank inbred mouse strains by predicted vulnerability or resilience to a disease state, supporting rational background selection for preclinical studies. The approach prioritizes among hundreds of inbred strains spanning common inbreds, recombinant inbreds, Collaborative Cross (CC) strains, wild-derived strains, and select virtual F1 hybrids (computationally simulated strain crosses). Inclusion of hybrids, particularly for C57BL/6 strains provides recommendations suitable for integrating existing gene deletion and other engineered dominant mutations into new models. Here we present the Strain Recommender approach and demonstrate its utility for rational model selection across a range of disease contexts.

Using two complimentary study designs, we validated the approach prospectively in a connective tissue disorder study using CC strains and cardiovascular phenotyping, and we assessed broader performance through retrospective benchmarking against curated background-dependent effects reported in the Mouse Genome Database (Eppig et al., 2015). Together, these analyses demonstrate that integrating disease-associated gene signatures with strain-specific transcriptome predictions effectively prioritizes background selection and broadens the range of strain backgrounds in preclinical research.

## RESULTS

### Overview of the Strain Recommender Workflow and Outputs

Strain Recommender prioritizes mouse strain backgrounds (and virtual C57BL/6J F1s) for disease research by ranking strains according to the similarity between their transcriptional profiles and disease-associated gene sets, prioritizing strains most likely to recapitulate the molecular features of a disease state (Fig. 1a). The approach is grounded in the premise that strain-specific differences in disease-related molecular networks influence tolerance to deleterious genetic and environmental perturbations, resulting in differential vulnerability to disease (Fig. 1b).

**Fig 1.**
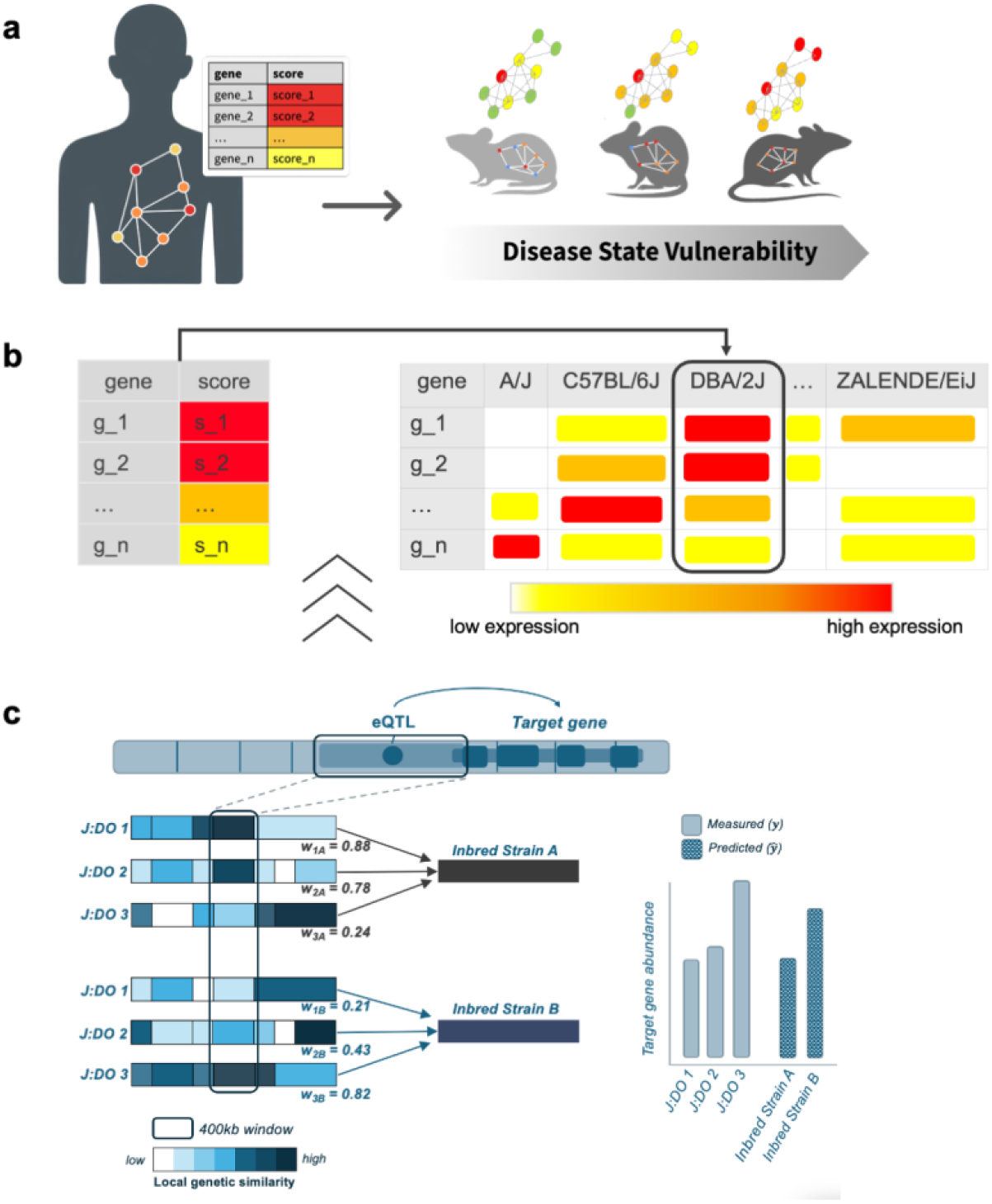
Strain Recommender Approach. **a**, Strain Recommender predicts which mouse strains best resemble a disease state. Strains less tolerant to deleterious mutations (red node) exhibit greater disease severity and closer resemblance to the disease. **b**, Resemblance to the disease state is quantified by comparing a gene score, capturing the direction and magnitude of effect (e.g., log2FC), to the imputed gene expression profile of each strain. Expression matrices are organized with genes in rows and strains (e.g., A/J, C57BL/6J, DBA/2J) in columns. Here, DBA/2J is predicted to most closely resemble the disease state. Colors indicate higher (red) or lower (yellow) relative expression. **c**, Transcriptomes are imputed for each strain by assigning a weight (*w*_*ij*_) based on the local genetic similarity to the inbred strain in the 400 kilobase (kb) region surrounding the expression influencing (eQTL) variant. Color intensity reflects the degree of genetic similarity between the J:DO and the inbred strain in the 400 kb region. Predicted expression for each strain is a function of observed J:DO expression and their local genetic similarity to the inbred strain.

Operationally, the approach evaluates a disease state represented as a genomic signature—a weighted gene set in which each gene is assigned a signed quantitative score representing the direction and magnitude of association with disease—and compares this signature to a strain-specific gene expression resource to quantify molecular resemblance. Strains are ranked by a vulnerability score (Spearman correlation between gene scores and predicted strain expression), and an uncertainty estimate is returned for each strain using a permutation-derived false positive rate (FPR; Fig. 1c; Table 1).

**Table 1.** Age-related macular degeneration: a case study.

| Strain | Score | FPR |
| --- | --- | --- |
| PL/J | 0.72 | 0.018 |
| C3H/HeH | 0.56 | 0.046 |
| C3H/HeJ | 0.55 | 0.048 |
| ... | ... | ... |
| BXD162 | -0.55 | 0.047 |
| LEWES/EiJ | -0.65 | 0.027 |
| WSB/EiJ | -0.81 | 0.008 |

Strain Recommender is available at https://recommender.jax.org/. The web-based tool allows users to input a disease of interest, select a related gene set from GeneWeaver (https://geneweaver.org/), a database of over 300,000 curated gene sets (Baker et al., 2012), and receive a ranked list of recommended mouse strains with associated confidence scores.

### Case Study: Prioritizing a Background for Age-Related Macular Degeneration

To illustrate use of the method on a familiar disease context, we applied Strain Recommender to age-related macular degeneration using two differential expression resources (peripheral retina and retinal pigment epithelium (Kim et al, 2018, GeneWeaver gene set GSID 412582). In this case, Strain Recommender identified PL/J, C3H/HeH, and C3H/HeJ as comparatively vulnerable among 657 mouse strains (Table 1). This prioritization is consistent with known strain-specific effects that interact with deleterious naturally occurring mutations such as *Pde6b<rd1>*, which was not included in the input gene set. These mutations are known to have greater effects in the PL/J, C3H/HeH and C3H/HeJ backgrounds (Chang et al, 2002; Table 1).

### Prospective Validation: Connective Tissue Disorder Strain Prioritization and Phenotyping

We next evaluated whether Strain Recommender can prospectively anticipate disease-relevant phenotypic differences between strains. We focused on connective tissue disorders (CTD) that affect musculoskeletal tissues, vasculature and cardiac structure, and constructed a genomic representation of CTD using Marfan syndrome-associated gene sets derived from differential gene expression in thoracic aortic tissue (GeneWeaver gene set GSID 412566; Hansen et al., 2019). Strain Recommender predicted CC036/Unc to be more vulnerable to CTD and CC027/GeniUnc to be relatively resilient with vulnerability scores 0.202 (FPR = 0.297) and - 0.347 (FPR = 0.177), respectively (Supplementary Table 1). Consistent with these predictions, multivariate analysis of electrocardiography and echocardiography measurements, known to be affected in CTD (Milewicz et al., 2005; De Backer et al., 2006; de Witte et al., 2011; The Marfan Foundation), revealed cardiovascular differences consistent with disease that were significantly greater in CC036/UncJ than in CC027/GeniUncJ, as measured by electrocardiography (*F*(7,33) = 8.08, *p* < 0.001) and echocardiography (*F*(11, 28) = 3.17, *p* = 0.003) (Supplementary Methods S2; Supplementary Tables 2 and 3). These results demonstrate that Strain Recommender prospectively predicted disease-relevant phenotypic differences between strains, suggesting that introducing a disease-relevant perturbation would be more likely to provoke a detectable response in CC036/UncJ than in CC027/GeniUncJ.

### Global Retrospective Validation of Background Effects

To assess performance across a broad range of diseases, we conducted a global retrospective validation using curated reports of strain-dependent background effects from the Mouse Genome Database (MGD; Baldarelli, et al., 2024). The dataset included 898 curated strain comparisons drawn from the published literature and annotated Mammalian Phenotype (MP) Ontology terms (Bello et al, 2025), of which 772 involved strains represented in Strain Recommender. MP terms were mapped to disease via the Human Phenotype Ontology (see Methods). Disease-relevant gene sets were programmatically retrieved from GeneWeaver’s database (https://geneweaver.org/, accessed September 10, 2024), including many obtained from transcriptome data repositories such as Short Read Archive and Gene Expression Omnibus, providing coverage for 6,488 of 8,856 diseases (73.3%), of which 5,890 (90.8%) met the FPR ≤ 0.05 threshold and were included in the analysis (Supplementary Tables 4, 5).

The analysis consisted of 2,014 distinct sets of comparisons, each defined by unique combinations of gene set, study, MP term, and strain-pair, applied combinatorially to evaluate disease-vulnerability predictions. For each disease, predicted relative vulnerability between strain-pairs was compared to experimentally observed outcomes in the curated MGD dataset (Fig. 2; Supplementary Table 4), scored in binary terms as correct (1) or incorrect (0). Performance was quantified as per-disease sensitivity, defined as the proportion of correct strain-pair predictions for a given disease, and summarized as mean sensitivity across diseases (Fig. 2b).

**Fig 2.**
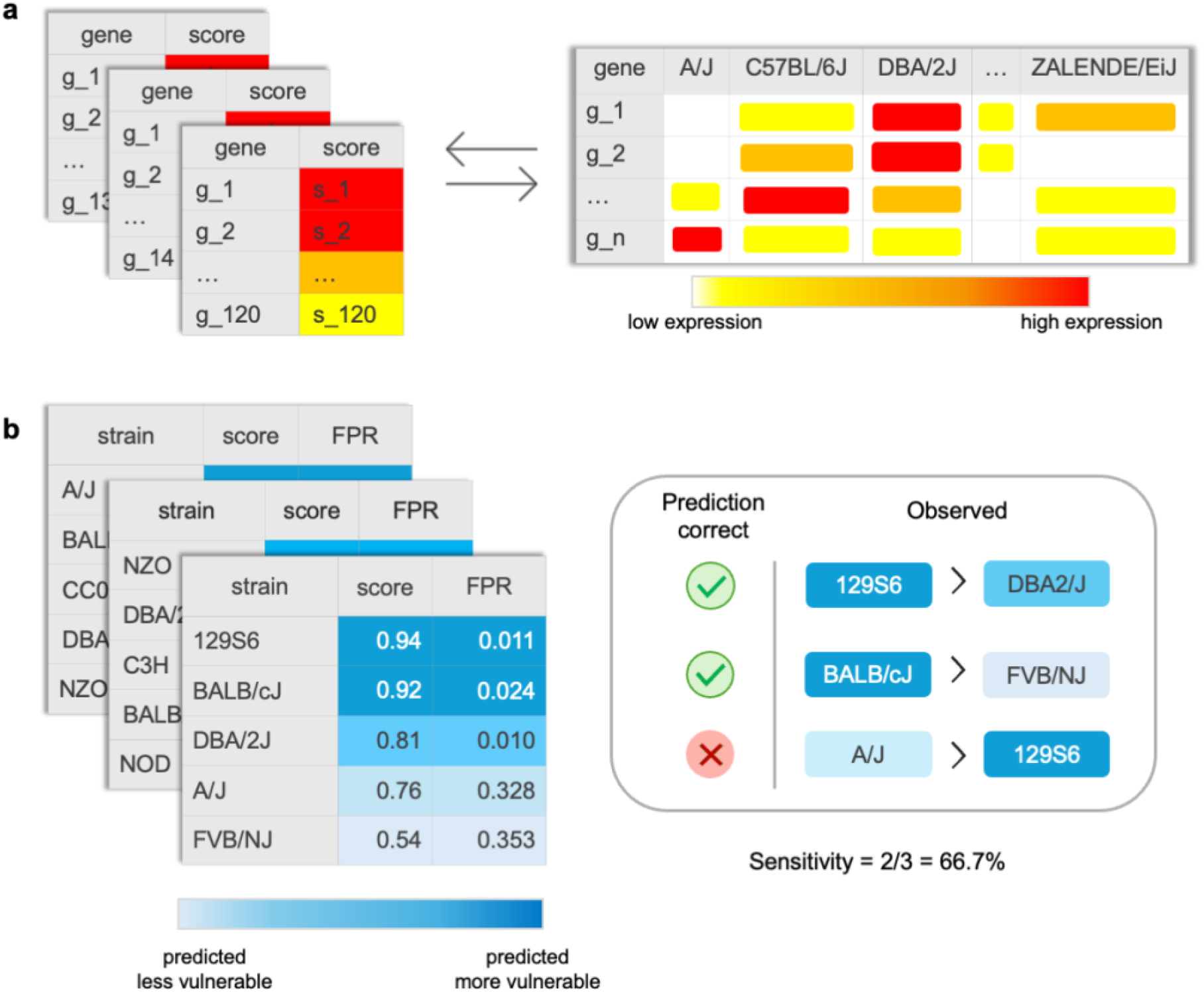
Global validation analysis approach. **a**, Disease-relevant gene sets extracted from the GeneWeaver database were input to Strain Recommender and compared to mESC imputed transcriptomes of over 600 inbred mouse strains. **b**, For each gene set used as input, Strain Recommender output a predicted strain vulnerability score and estimated FPR. Sensitivity (true positive rate) was then calculated per disease across all comparosons. In this example, Strain Recommender correctly predicted 2 of 3 (66.7%) strain comparisons in the validation set, illustrated by the mouse pairings, i.e., Strain Recommender predicted 129S6 with score 0.94 as more vulnerable than DBA/2J with score 0.81, which matches the observed strain ranks in the validation set (right).

Across all diseases, Strain Recommender achieved a mean overall sensitivity of 94.4% (95% CI: 94.0 – 94.8%), and 92.3% (95% CI: 91.6 – 93.0%) across rare diseases. In absolute terms, Strain Recommender achieved sensitivity > 50% for 5,578 of 5,890 diseases (94.7%) and 2,507 of 2,598 rare diseases (96.5%). When stratified by per-disease sensitivity, Strain Recommender achieved high sensitivity (≥90%) for 86.6% of all diseases and 81.5% of rare diseases, moderate sensitivity for 4.7% and 6.5%, respectively, low sensitivity for 3.4% and 8.5%, respectively, and poor sensitivity (≤50%) for 5.3% and 3.5%, respectively (Fig. 3).

**Fig 3.**
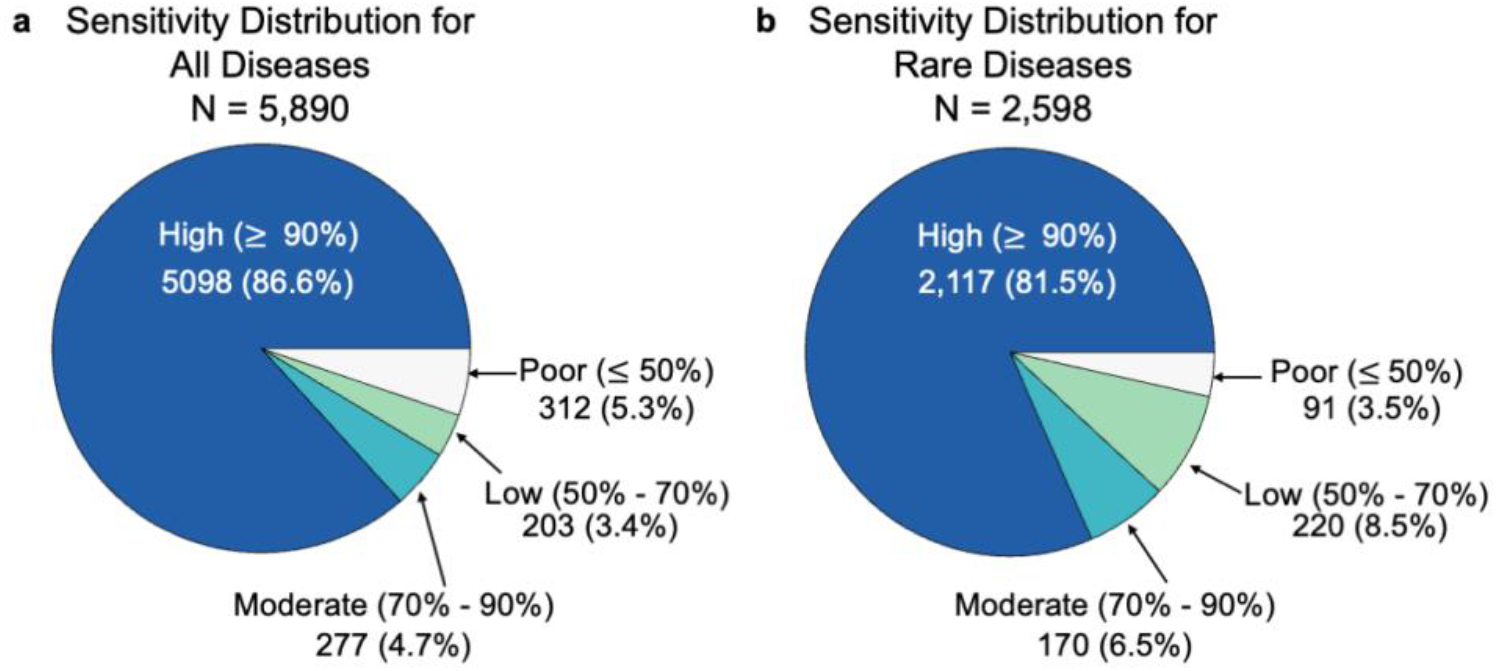
Strain Recommender sensitivity distribution across diseases. **a**, Strain Recommender makes the correct prediction 94.7% of the time across all diseases (N = 5,890), and **b**, and 96.5% of the time across rare diseases (N = 2,598). Sensitivity is the proportion of correct predictions per disease.

### Sensitivity Analyses to Identify Factors Associated with Performance

To identify factors associated with prediction performance, we conducted a series of sensitivity analyses examining gene set characteristics, FPR threshold, and dataset redundancy. The effect of gene set size, provenance and species were assessed after accounting for random effects of gene set and study. Moderately sized gene sets (50-99 genes) yielded significantly better performance than other size categories (gene sets of size < 50, 100-499, and 500+; 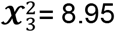, *p* = 0.030). Species and expert curation status did not significantly affect performance after accounting for gene set size (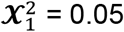, *p* = 0.820 and 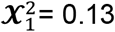, *p* = 0.721, respectively).

Relaxing the FPR threshold from 0.05 to 0.10 and 0.15 resulted in mean sensitivity dropping from 94.4% (95% CI: 94.0-94.8%) to 86.6% (95% CI: 86.0-87.2%) and 83.8% (95% CI: 83.2-84.4%), respectively, with a corresponding increase in disease coverage from 5,890 to 6,039 and 6,135, respectively. At an FPR threshold of 0.20, sensitivity dropped substantially to 46.1% (95% CI: 45.7-46.5%), suggesting 0.05 as a reasonable default threshold (Supplementary Table 4).

We estimated validity over the widest possible range of diseases to reflect expected outcomes from actual usage of the tool; however, multiple diseases are associated with a similar combination of ontology annotations. To assess sensitivity on non-redundant comparisons, disease-specific sensitivities and overall sensitivity were also calculated over a subset of 2,014 diseases in which each disease-specific sensitivity was based on a unique set of comparisons, providing a conservative estimate of real-world performance. Mean sensitivity was 87.0% (95% CI: 86.0-88.0%) with a median of 100%. As in the primary analysis, high, moderate, low, and poor sensitivity were defined as ≥90%, 70-90%, 50-70% and ≤50%, respectively (Fig. 3). Strain Recommender achieved high sensitivity for 1,374 (68.2%) diseases, moderate sensitivity for 237 (11.8%) diseases, low sensitivity for 168 (8.3%) diseases, and poor sensitivity for 235 (11.7%) diseases (Supplementary Table 4).

### Improving Existing Mouse Models by Identifying Alternative Backgrounds

Across nearly all diseases evaluated, Strain Recommender identified strains predicted to more closely resemble the disease state than background strains previously reported in the literature. Specifically, in over 88% of diseases, at least one strain with more extreme predicted vulnerability was identified relative to the most extreme strain reported in the MGD (Fig. 4). Notably, 185 inbred strains (28.2%) were predicted to resemble disease states more frequently than C57BL/6J, the most common background for modeling human disease (Fig. 4; Supplementary Table 6), highlighting substantial opportunities to refine background selection and improve model relevance.

**Fig 4.**
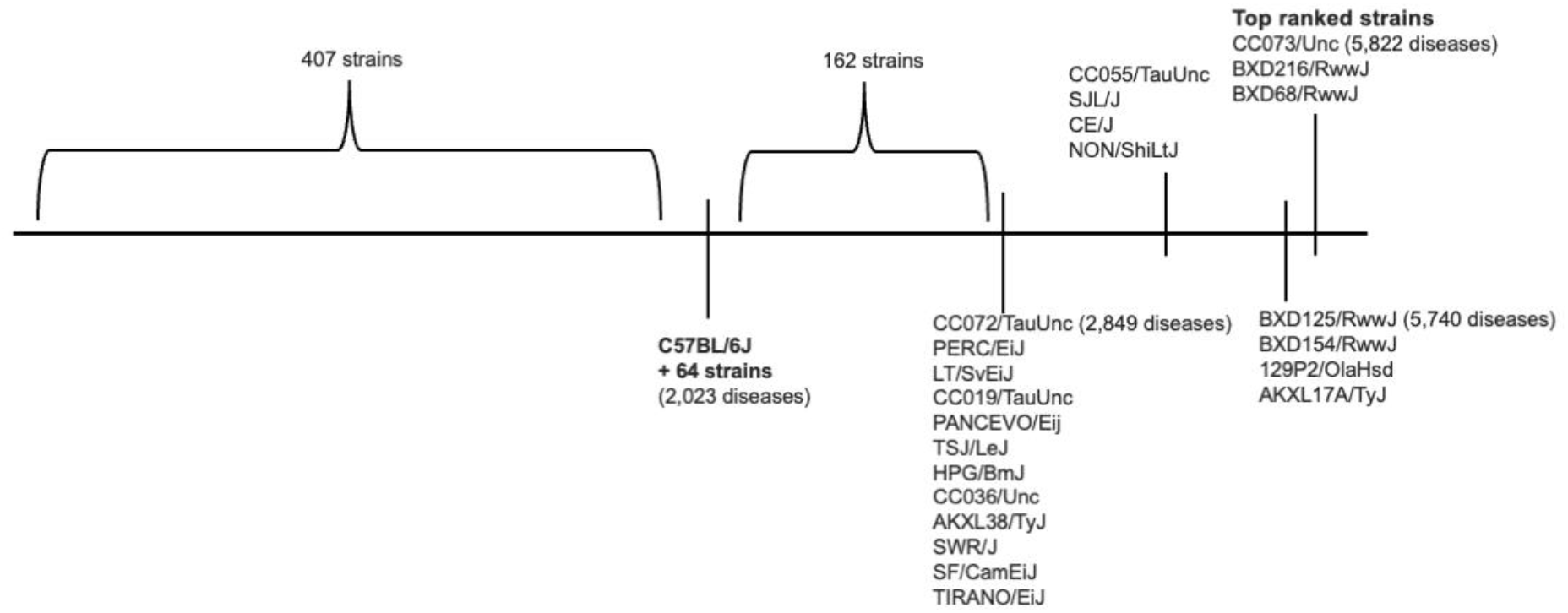
Strain Recommender identifies a strain with more extreme predicted vulnerability than previously tested models for 88% of diseases. The vulnerability landscape of the 657 inbred strains across 6,488 diseases illustrates that a strain with more extreme predicted vulnerability is identified by Strain Recommender than what is reported in MGD over 88% of the time (Supplementary Table 5). The figure illustrates the ranking of strains based on the number of times the strain was predicted to be vulnerable to a disease at FPR ≤ 0.05 (lower to higher rank from left to right).

## DISCUSSION

Genetic background is a critical yet often underappreciated determinant of model validity in laboratory mice and other experimental systems. Strain-specific differences in molecular networks can modulate the penetrance and expressivity of disease-associated perturbations, influencing both phenotype severity and reproducibility across studies. Strain Recommender addresses this longstanding challenge by providing a data-driven framework to prioritize strain backgrounds that are predicted to be relatively vulnerable or resilient to a user-defined disease state, enabling more rational selection of backgrounds before investing in time- and resource-intensive model development.

Strain Recommender integrates curated disease-relevant gene sets with imputed, strain-specific transcriptomic profiles to rank strains by the concordance between the disease-associated molecular signature and each strain’s predicted transcriptional state. This operationalizes a simple, testable hypothesis: strains whose baseline transcriptional state more closely resembles a disease signature may be less tolerant to additional perturbations, e.g., engineered mutations or environmental stressors (Nadeau, 2001) and therefore more likely to manifest disease-relevant endophenotypes (Gould & Gottesman, 2006).

In both prospective and retrospective evaluations, Strain Recommender demonstrated strong performance and broad coverage across diseases, including rare diseases. Importantly, the platform frequently identified alternative strain backgrounds predicted to more strongly resemble the disease state than the background reported in the literature (Fig. 4; Bhatt et al. 2020; Shineman et al., 2011; Long et al., 2021; Howe et al., 2018), highlighting opportunities to improve existing models and reduce reliance on historically favored strains that may be suboptimal for specific disease contexts.

Strain Recommender’s transcriptome imputation shares conceptual roots with approaches developed for human genetics (e.g., PrediXcan [Gamazon et al., 2015] and related methods) but differs in its construction and intended use. Rather than predicting gene expression from variant-level features or raw genomic sequence alone (Dubey and Shen, 2026), Strain Recommender predicts expression in each inbred strain from a weighted combination of measured expression across a sample of Diversity Outbred (J:DO) mice, which harbors nearly 90% of the segregating variation observed in widely studied inbred mouse strains (Roberts et al, 2007). Weights reflect local genetic similarity between each J:DO individual and the target strain in the regulatory region of each gene, capturing the combined effects of linked regulatory variants within each gene’s *cis*-regulatory network. This design leverages two advantages of mouse resources: fixed, well-characterized inbred genomes that support reproducible similarity calculations, and dense genome-wide genotype data across hundreds of strains (GenomeMUSter; Ball et al., 2024; RRID:SCR_024214). Importantly, imputed gene expression is not the end goal; it provides a scalable strain-specific molecular substrate that enables the vulnerability score and reflects the extent to which a strain’s transcriptional state molecularly resembles a disease state, and therefore its likely capacity to buffer or succumb to additional deleterious genetic variation. This distinction matters for interpretation: Strain Recommender is intended to prioritize standardized, fully reproducible backgrounds for model development, not to assert that the imputed transcriptome fully captures all aspects of disease-relevant expression across tissues, developmental stages or environmental context.

### Generalizability and Practical Use

The current implementation focuses on inbred mouse strains and isogenic hybrids. Not all of the recommended strains will be readily available, and some present challenges in breeding and other reproductive characteristics, however, users can select from a number of highly ranked strains based on practical considerations. Engineering on poorly characterized genetic backgrounds is not without challenges, but CRISPR/cas9 and recent work to generate diverse strain embryonic stem cells has greatly expanded the number of strains readily available for engineering (Czechanski et al., 2014). The inclusion of virtual F1 hybrids presents an opportunity to improve existing disease models carrying mutations on a C57BL/6 background, such as widely used knock-out mouse resources. A simple intercross with a predicted vulnerable strain could yield a more disease-relevant model without the time and expense of engineering the mutation onto an alternative strain background.

The underlying recommender framework can be adapted to other model organisms where comparable genotype resources and expression training sets exist (e.g. rats [Johnson et al., 2023] and *Drosophila* [Mackay et al., 2013]). In addition, the approach could be applied to genetically defined cellular resources (Pantazis et al., 2022) to bridge *in vitro* and *in vivo* systems. By enabling earlier, data-driven strain prioritization, Strain Recommender can reduce the cost and animal use associated with exploratory strain screening and improve reproducibility by motivating explicit, evidence-based background selection. This aligns with the NIH’s 3Rs principles—Replacement, Reduction, and Refinement (Russell and Burch 1959; Strech and Dirnagl, 2019), while recognizing that for some applications, in *vivo* systems are indispensable.

### Limitations and Future Directions

Several limitations should be acknowledged. First, the accuracy of imputed transcriptomes depends on the quality and resolution of genotype resources and gene regulatory mappings, and performance can vary across gene sets and disease contexts. Improved profiling of widely used strains will improve the precision and accuracy of predictions. Second, approximately 16% of named diseases are not represented in curated datasets. For these, input gene sets can be constructed from new molecular profiling, studies of similar diseases, gene-trait associations for characteristics of the disease and many other approaches. For extremely rare diseases and *de novo* mutations, these strategies are essential. Finally, high aggregate sensitivity can mask variable performance across diseases, especially for insufficiently characterized conditions with limited gene set support. Future work could expand disease-state representations beyond gene sets and employ other algorithmic approaches to infer disease similarity and improve transferability across related phenotypes.

### Outlook

Although Strain Recommender is validated using curated mutation-by-background interactions, its utility extends beyond monogenic models. The framework is well-suited for prioritizing backgrounds for complex disease modeling, where multiple variants and regulatory networks jointly shape susceptibility. By supporting rational, evidence-based strain selection, Strain Recommender has the potential to improve the fidelity of research in experimental disease models and strengthen the translational value of preclinical studies.

## METHODS

### Strain Recommender Workflow Overview

Strain Recommender prioritizes mouse strain backgrounds by quantifying concordance between (i) a genomic representation of a disease state with signed quantitative gene scores and (ii) strain-specific transcriptomic profiles imputed for a panel of 657 inbred strains in mouse embryonic stem cells (mESC). For each strain, the method returns a disease vulnerability score and an uncertainty estimate (false positive rate, FPR) derived from a permutation-based null model.

#### Disease-state input (gene set definition)

Strain Recommender requires a genomic quantification of the disease state consisting of gene identifiers and gene scores indicating the direction and magnitude of association with disease (for example, signed differential-expression statistics). The gene set is harmonized by (i) translating gene identifiers to orthologous mouse Ensembl gene identifiers through GeneWeaver’s AON API (https://geneweaver.jax.org/aon/api/docs), (ii) resolving multiple gene scores for the same gene by retaining only the highest absolute score, and (iii) filtering the gene set to include only genes in the strain expression resource. The number of genes (*n*) used to predict strain vulnerability is tracked because it parameterizes the uncertainty model (below).

#### Strain expression resource

A key challenge for strain prioritization is the absence of comprehensive transcriptomic profiling across large strain panels under relevant disease contexts. Strain Recommender addresses this gap through transcriptome imputation across 657 strains using gene expression data from mESC. Machine learning models were developed on mESC data sampled from 185 genomically distinct diversity outbred (J:DO) mice (Skelly et al, 2020; Fig. 1c). The imputed expression matrix is used as a fixed input resource for all downstream scoring.

Gene-specific Gaussian ridge regression models were trained using 5-fold cross-validation, implemented via the R package *glmnet* (Friedman et al, 2010). Rather than using variant state directly as predictors, each gene model predicts strain expression as a weighted combination of observed J:DO expression profiles. Each gene model incorporates local expression quantitative trait loci (eQTL) information and regional haplotype structure. Weights are constructed from genetic distances within a 400 kilobase window around the variant influencing expression, upweighting expression profiles of genetically similar J:DO individuals (Fig. 1c).

The fitted models were then applied to 657 inbred strains, using GenomeMUSter genomes to compute genetic similarity and corresponding weights between J:DO animals and inbred strains. Full model specification and the local-weighting construction are provided in Supplementary Methods S1.

#### Strain vulnerability score

For each strain, Strain Recommender computes a vulnerability score as the Spearman’s rank correlation (*ρ*) between the strain’s imputed expression values (*n* genes) and the disease gene scores for those same genes. Scores range from -1 to +1. Positive values indicate that genes up-scored in the disease state tend to be more highly expressed in that strain (and genes down-scored in disease tend to be lower), consistent with greater molecular resemblance (“vulnerable”); negative values indicate the opposite pattern (“resilient”).

#### Uncertainty quantification via a permutation-based False Positive Rate (FPR)

To quantify uncertainty in each strain’s vulnerability score, we estimate a one-sided FPR with a permutation test that generates a null distribution specific to the input gene set size (*n*), accounting for uncertainty in both the gene set input and the imputed expression resource. For each strain, 4,000 null scores are generated by: (1) selecting a random strain from the imputed transcriptome resource; (2) randomly sampling *n* genes from that random strain; and (3) computing Spearman’s *ρ* between the sampled expression values and the observed disease gene scores. The one-sided FPR is computed as the fraction of null correlations at least as extreme as the observed correlation in the direction of the observed effect (i.e., right tail for positive “vulnerable” scores; left tail for negative “resilient” scores).

### Strain Recommender validation

We evaluated Strain Recommender prospectively in a connective tissue disorder (CTD) experiment and retrospectively at scale using curated background-dependent effects from the Mouse Genome Database (MGD). The age-related macular degeneration example is treated as a case study and is therefore described in Results; Methods will reference the same scoring workflow above without re-defining it.

#### Prospective validation: connective tissue disorders

The CTD gene set construction and strain selection are described in Results. Electrocardiography was performed for 12 female and 10 male CC036/UncJ mice and 12 female and 7 male CC027/GeniUncJ mice (41 measures). Echocardiography was performed for 11 female and 10 male CC036/UncJ mice and 12 female and 7 male CC027/GeniUncJ mice (40 measures). Based on literature review, 7 electrocardiograph and 11 echocardiograph measures most relevant to Marfan Syndrome were analyzed using MANOVA to test for an overall strain effect across measures (Supplementary Table 2). Follow up ANOVA models were run for each of the 18 measures, with Benjamini-Hochberg correction for multiple comparisons and a significance level of 0.05 (Supplementary Table 3; Supplementary Methods S2).

#### Global retrospective validation: ground truth dataset

To retrospectively validate predictions, we used a curated dataset from MGD of observed variable strain background effects. Each entry includes a study identifier, two strain names, and an annotation indicating which strain was experimentally observed to be more phenotypically similar (i.e., more vulnerable) relative to the MP term. Dataset curation and filtering criteria are described in the Results.

#### Ontology mapping from Mammalian Phenotype (MP) terms to human disease terms

Because evaluation is summarized at the level of human disease, MP terms in the curated validation dataset were mapped to human disease terms using publicly available ontology databases and services so that each disease in the validation set is traceable to published strain background studies. Specifically, we used the JAX Ontology Service (https://ontology.jax.org/, accessed September 4, 2024) and Unified Phenotype Ontology (uPheno; https://www.ebi.ac.uk/ols4/ontologies/upheno, accessed September 3, 2024) to map MP terms to HP terms in the Human Phenotype Ontology (HPO) (Gargano et al, 2024), which was also used to map HP terms to human disease (https://hpo.jax.org/data/annotations, accessed September 27, 2024, Supplementary Table 7). Rare diseases were identified using ORPHAcodes downloaded from Orphanet Reports (https://www.orpha.net/en/other-information/reports-List of rare diseases and synonyms in alphabetical order accessed October 25, 2024).

#### Genomic quantification of disease state for retrospective evaluation

To assemble disease-relevant gene sets for retrospective validation, MP terms and descriptions, mapped HPO terms and descriptions, and disease labels were queried via the GeneWeaver Search AON API (https://geneweaver.jax.org/api/genesets/search, accessed September 10, 2024), returning candidate gene sets for each disease context. Public gene sets were included if they (i) contained at least 3 genes, (ii) had continuous gene scores that varied across genes, and (iii) were derived from human or mouse studies (Supplementary Table 5).

#### Performance metrics and statistical analysis

Retrospective evaluation focuses on predictions with low uncertainty; specifically, strain-pair comparisons were included when relevant strain scores met FPR ≤ 0.05. This filtering yielded 58 gene sets used for the primary retrospective analysis. If the highest-ranked strain in an observed study did not meet the FPR threshold, the gene set was deemed insufficient to support a confident prediction for that strain in retrospective benchmarking.

Predictions were scored as correct (true positive = 1) if the predicted strain vulnerabilities matched observed strain background effects found in the literature (MGD), and incorrect (false negative = 0) otherwise. Disease-specific sensitivity was defined as the number of true positives divided by the number of cases (true positives + false negatives) within disease. Mean overall sensitivity was calculated as the unweighted mean of disease-specific sensitivities (macro-average), with 95% confidence intervals based on the Normal distribution, justified by the Central Limit Theorem (n = 5,890 diseases). All computations were performed in R (R Core Team, 2025).

#### Sensitivity analyses

Fixed effects of species, gene set size, and curator review status were tested with likelihood ratio tests on nested logistic mixed-effects models, with random effects for gene set and study identifier. Models were fit using the *glmer* function in the *lme4* package (Bates et al., 2015). To evaluate the sensitivity of results to the FPR threshold, disease-specific and mean overall sensitivities were calculated across a range of FPR thresholds using the complete dataset prior to threshold filtering (Supplementary Table 4). A non-redundant subset of diseases was constructed by identifying diseases sharing identical sets of comparisons and retaining a single representative disease per unique comparison set (*n* = 2,014), with the first listed disease used as the representative.

## Acknowledgements

This work was supported by The Jackson Laboratory’s Center for Precision Genetics NIH U54OD030187 and The Jackson Laboratory. We gratefully acknowledge the contribution of JAX Data Science including John Bluis, Sejal Desai, Ben Walton, Beena Kadakkuzha, Georgi Kolishovski, Sneta Kayastha, Hao He, Sara Davis, Dave Walton, and Vivek M. Philip with support from NCI CCSG P30CA034196. We thank Patsy Nishina and Martin Pera for sharing their expertise on age-related macular degeneration and retinal degeneration in mice. We thank Leona Gagnon and Kathy Snow for technical management of the CTD prospective validation study, and the JAX Center for Biometric Analysis for cardiac phenotyping, with support from NCI CCSG P30CA034196. MGD and the Mammalian Phenotype Ontology are funded by NIH U24HG000330. We greatly appreciate the editorial support from Miguel Skirzewski of JAX Research Program Development.

## Contributions

R.L.B., P.N.R. and E.J.C. conceptualized the project for the JAX Center for Precision Genetics under the leadership of S.A.M. R.L.B. developed the transcriptome imputation models. M.J.K. and R.L.B. trained the models and predicted mESC transcriptomes for inbred strains. M.W.G. created the eQTL and gene expression data services for transcriptome imputation. G.M. and G.K. analyzed raw data from SRA and GEO and curated additional disease-relevant gene sets into GeneWeaver. The retrospective validation dataset was assembled by M.L.B. and C.L.S. under the leadership of C.J.B., annotated by M.L.B., and made computable by H.S.F., T.N-M., A.K., and R.L.B. M.A.G., M.J.K., and A.K. created the ontology map of MP, HP, and disease terms. F.C., A.K.B-L., and A.K. extracted disease-relevant gene sets for the retrospective validation from GeneWeaver. A.K. and R.L.B. evaluated Strain Recommender performance on the prospective and retrospective validation sets and deposited the codebase in GitHub. A.K., R.L.B., and E.J.C. wrote the paper with input from all authors.

